# Creation of gene expression database on preeclampsia-affected human placenta

**DOI:** 10.1101/102012

**Authors:** Oleksandr Lykhenko, Alina Frolova, Maria Obolenska

## Abstract

Publication of gene expression raw data in open access at online resources like NCBI or ArrayExpress made it possible to use these data for cross-experiment integrative analysis and make new insights into biological phenomena. However, most popular of the present online resources are meant to be archives rather than ready for immediate access and interpretation databases. Data uploaded by independent contributors is not standardized and sometimes incomplete and needs further processing before it is ready for the analysis. Hence, the need for a specialized database appears.

Given in this article is the description of the database that was created after processing a collection of 33 relevant datasets on pre-eclampsia-affected human placenta. Data processing includes the choice of relevant experiments from ArrayExpress database, the experiment sample attributes standardization according to MeSH term dictionary and Experimental Factor Ontology and the completion of missing data using information from the corresponding articles and authors.

A database of more than 1000 samples contains sufficient sample-wise metadata for them to be arranged into relevant case-control groups. Metadata includes information on biological specimen, donor’s diagnosis, gestational age, mode of delivery etc. The average size of these groups will be higher than it is in separate experiments. This will reduce experiment bias and enhance statistical accuracy of the subsequent analysis such as search for differentially expressed genes or inferring gene networks. The article concludes with the guidelines for the microarray experiment metadata uploading for future contributors.

## Background

Open gene expression databases have over time acquired tremendous amounts of data [9, 16]. It is now possible to make original discoveries just by analyzing these data. An *integrative analysis* is one of the approaches in this case. Unlike *meta-analysis*, which is essentially adding up results of different experiments, an integrative analysis implies merging or integration of raw data which, as studies show [11, 12], identifies significantly more differentially expressed genes [19]. Besides, integrative analysis provides larger sample sets and thus increases statistical significance and reduces experimental bias [20] which makes it most useful in cases when individual experiments’ average sample set is small.

To identify which pieces of data are similar enough for the integration to make sense each individual sample must be supplemented with at least minimal data. Here and later these sample clinical and biological information will be called sample *metadata* (do not confuse with meta-analysis, a method to unite results of different studies). Unfortunately, most popular public databases like NCBI or ArrayExpress not always contain sufficient sample-wise metadata. There are several reasons for that. The main one is that the integration is not a primary goal for these databases. As is stated in [17] ArrayExpress primary goal is to serve as an archive for microarray data associated with scientific publications and other research. Also, since the data was uploaded by independent contributors the biological sample metadata is not standardized and, consequently, not ready for immediate automated access by a search query. The choice of relevant characteristics of biological sample is also up to a contributor and those characteristics do not always match the needed ones for the data integration.

To address these problems investigators develop specialized databases. Here we present a curated collection of publicly available datasets on preeclampsia-affected human placenta.

Pre-eclampsia [1] is a disorder that occurs only during pregnancy and the post-partum period and affects both the mother and the unborn baby. Affecting at least 5-8% of all pregnancies, it is a rapidly progressive condition characterized by high blood pressure and the presence of protein in the urine. Typically, pre-eclampsia occurs after 20 weeks gestation (in the late 2nd or 3rd trimesters or middle to late pregnancy) and up to six weeks postpartum (after delivery), though in rare cases it can occur earlier than 20 weeks. Globally, pre-eclampsia and other hypertensive disorders of pregnancy are a leading cause of maternal and infant illness and death.

Although etiology and pathogenesis of pre-eclampsia are still unknown numerous studies (listed in Supplement 1) point at the gene disregulation in placenta to be one of the possible prerequisites for the disease. Besides, there are also known cases when placenta develops without fetus, called molar pregnancy [3], which are associated with very early-onset pre-eclampsia. This is why we focused our study on gene expression in placenta. Finally, we chose cDNA microarray technology as most popular in pre-eclampsia studies among ones providing information on the whole transcriptome.

Our database now contains sufficient metadata for the samples to be united into relevant case-control groups for the subsequent integrative analysis and further search for differentially expressed genes and inferring gene networks.

## Methods

Our software is written in Python language using Django framework for web interface development and Postgres for relational database support. All experiment and sample metadata were automatically extracted from ArrayExpress database via Bioservices which is a Python interface to ArrayExpress. NCBI database was used to supplement the missing data along with the corresponding scientific articles and authors personally.

Source code and database backup file are available at GitHub: https://github.com/Sashkow/placenta-preeclampsia

Web interface for our database at its current stage can be accessed at: http://194.44.31.241:24173/

## Results

The database at its current stage contains a total of 32 experiment datasets and more than 1000 placenta samples, about 900 of which contain minimal metadata for relevant cross-experiment study groups to be constructed, which is diagnosis, gestational age and tissue type. A separate group of 11 experiment datasets and 300 samples are in vitro experiments with cell cultures and immortalized cell lines as biological samples. Apart from pre-eclampsia, some related complication were considered including fetal growth retardation, HELLP syndrome and some genetic disorders such as mosaicism and trisomy of chromosome 16.

### Database structure

Figure 1 shows general structure of the database. Rectangles represent tables in database. Tables are linked with one-to-many and many-to-many relations. For example, Experiments table is in many-to-many relation with Microarrays table for each experiment may utilize multiple microarray platfoms and each platform can be used in multiple experiments. On the other hand, Experiments table is in one-to-many relation with Samples table as experiment may contain multiple samples while each sample can be a part of only *one* experiment. Experiments and Microarrays tables contain HStore field for storing mappings of strings to strings which are experiment/microarray attribute name-value pairs like “accession:E-GEOD-42424” in our case. SampleAttributes table containing all attributes for all samples is here to provide opportunity for different samples to have different attributes and to make new names reverse compatible with the originally downloaded ones. StandardSampleAttributeNames and StabdarSampleAttributeValues tables store standard terms for sample attribute names and values. The field named *additional_info* is HStore field with information about such as term source and short description. These tables for starndard terms also contain *synonyms* many-to-many fields with themselves providing lists of synonymous terms. Detailed explanation of the database design process can be found in Supplement 3.

**Figure 1:**
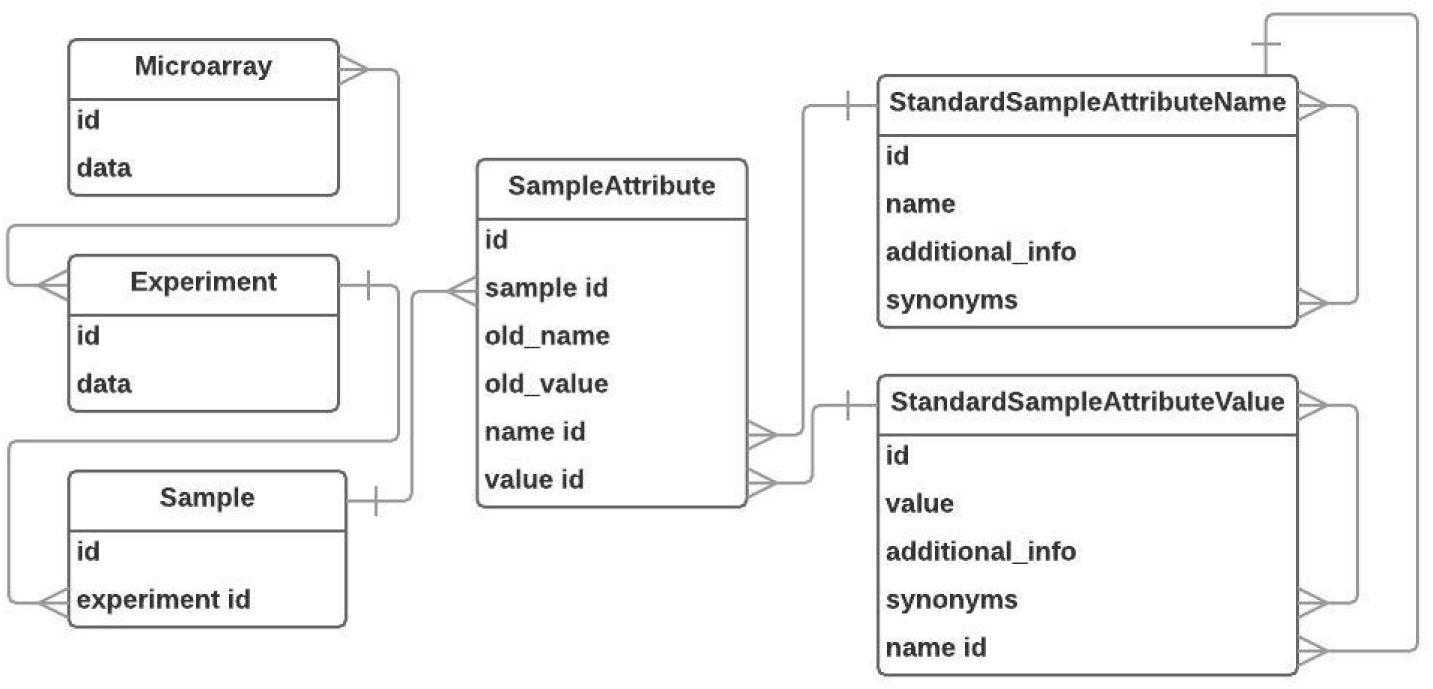
Database entity relation diagram.

**Figure 2:**
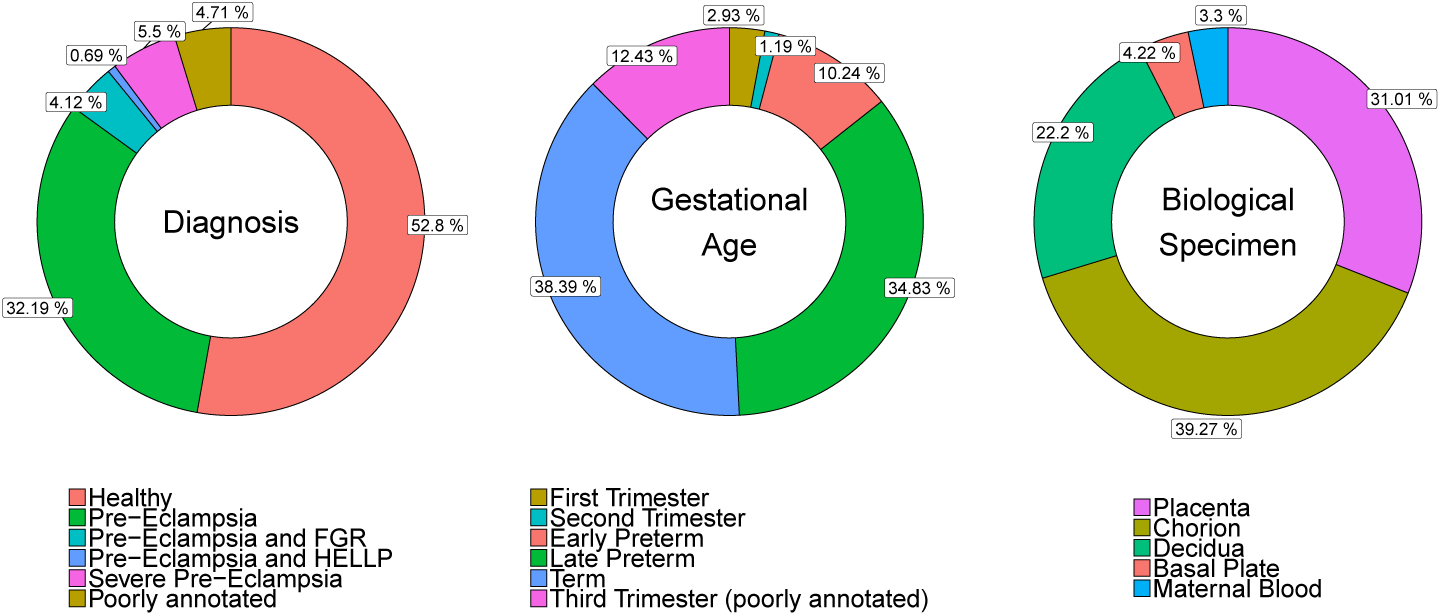
Sample core characteristics.

### Choice of datasets

The initial list of 43 relevant datasets was obtained as a result of ArrayExpress search by the following query: “preeclampsia OR pre-eclampsia OR preeclamptic OR pre-eclamptic” with results filtered by organism “Homo sapiens”, experiment type “rna assay”, experiment type “array assay”. E-GEOD-25906 was excluded due to data retrieval failure. E-MTAB-3732 was excluded since it is a compilation of microarray experiments for different diseases taken from publicly available sources. E-GEOD-15787, E-GEOD-22526, E-MEXP-1050 were excluded due to old microarray design or failure to find probe nucleodide sequences for the array. The full list of included and excluded can be found in Supplement 1.

### Sample attribute names and values standardization

Medical Subject Headings (MeSH) terms dictionary [4] was chosen as a standard for naming biological sample attribute names and values. We also used Ontology Lookup Service (OLS) as a secondary source in cases where no fitting MeSH term was found, since OLS utilizes multiple ontologies and has been recently updated to have more convenient interface than it used to. Furthermore, ArrayExpress itself uses one of OLS’s ontologies, namely Experimental Factor Ontology (EFO), to perform advanced search over genetic experiments and biological samples [5]. It means that our standardized samples metadata is potentially compatible with ArrayExpress and might be used to improve its search quality regarding datasets we have processed.

Here comes the list of standard sample attribute names and values, which are MeSH or other ontology terms, with lists of mappings onto sample attribute names and values originally downloaded from ArrayExpress. The format is the following:

~~~
Standard Name 1 (list of original names)
    Standard Value 1(list of original values)
    Standard Value2(list of original values)
    …
Standard Name 2(list of original names)
    Standard Value 1(list of original values)
    Standard Value 2(list of original values)
    …
…
~~~

Note that mapping is ambiguous meaning that original name may map onto different standard names depending on the context of the experiment. For example, original name “placenta” will map onto standard name “Chorion” if the context of the experiment suggests that. Some standard names do not map on any original name since that information was absent in ArrayExpress data and obtained manually either from the corresponding articles or from the authors personally. Here is a list of mappings for Diagnosis sample attribute provided as an example. The entire list of mappings can be found in Supplement 2.

~~~
Diagnosis ( diagnosis , subject status , disease , disease state
   , disease status , genotype , condition , group , phenotype ,
    classification , Disease State , Disease State )
Healthy ( normotensive , normotensive control patient ,
   normal , <empty>, healthy , control , Health , preterm
    labor , Normotensive control pregnancy , preterm )
Pre–Eclampsia (preeclamptic , pre–eclamptic patient ,
   preeclampsia , Pre–Eclampsia , EOPET , PE ,
   Preeclampsia after 20 weeks of gestation to 140/90
   mmHg , excretion of 0.3 g protein in 2 urine
   samples , early on set preeclampsia , late onset
preeclampsia , mild preeclampsia )
Pre–Eclampsia and HELLP
Pre–Eclampsia and FGR ( EOPET, pre–eclampsia and fgr ,
   pree clampsia=infant SGA)
Severe Preeclampsia ( severe preeclampsia , preeclampsia ,
    severe pre–eclampsia )
Fetal Growth Retardation ( IUGR , Fetal growth
   restriction (FGR) , infant SGA )
~~~

### Sample metadata completeness

While sample metadata from all considered datasets are now *standardized*, sample metadata remains far from complete. Yet it is already sufficient to tell whether any two of the samples are *comparable* i.e. whether they can be put to the same case-control group or not. We identified three key attributes that shall be our criteria for comparability: Diagnosis, Gestation Age, Biological Specimen - for only the samples of the same tissue at the same stage of the placental development under similar conditions can be put into the same study group whenever search for differentially expressed genes between control and deviation groups is to be performed.

**Gestational Age** is for some reason often not mentioned directly in sample data. However, it is possible to infer a meaningful category for gestational age by looking, for instance, at whether delivery was performed at term (37 to 41 week of gestation) or prematurely (20 to 37 weeks) or at pregnancy trimester if mentioned. Also, authors often publish gestation age mean and deviation for a study group which as well can be used to determine gestation age category for the samples. We used Gestational Age distribution for samples with known exact value to identify five gestation categories: first trimester, second trimester, early preterm, late preterm, term (Figure 3). Then we used approximate gestational age information for the samples with unknown exact Gestational Age value to put them into one of the five suggested gestation categories. Thus we determined gestation age for all samples to within gestational age category.

**Figure 3:**
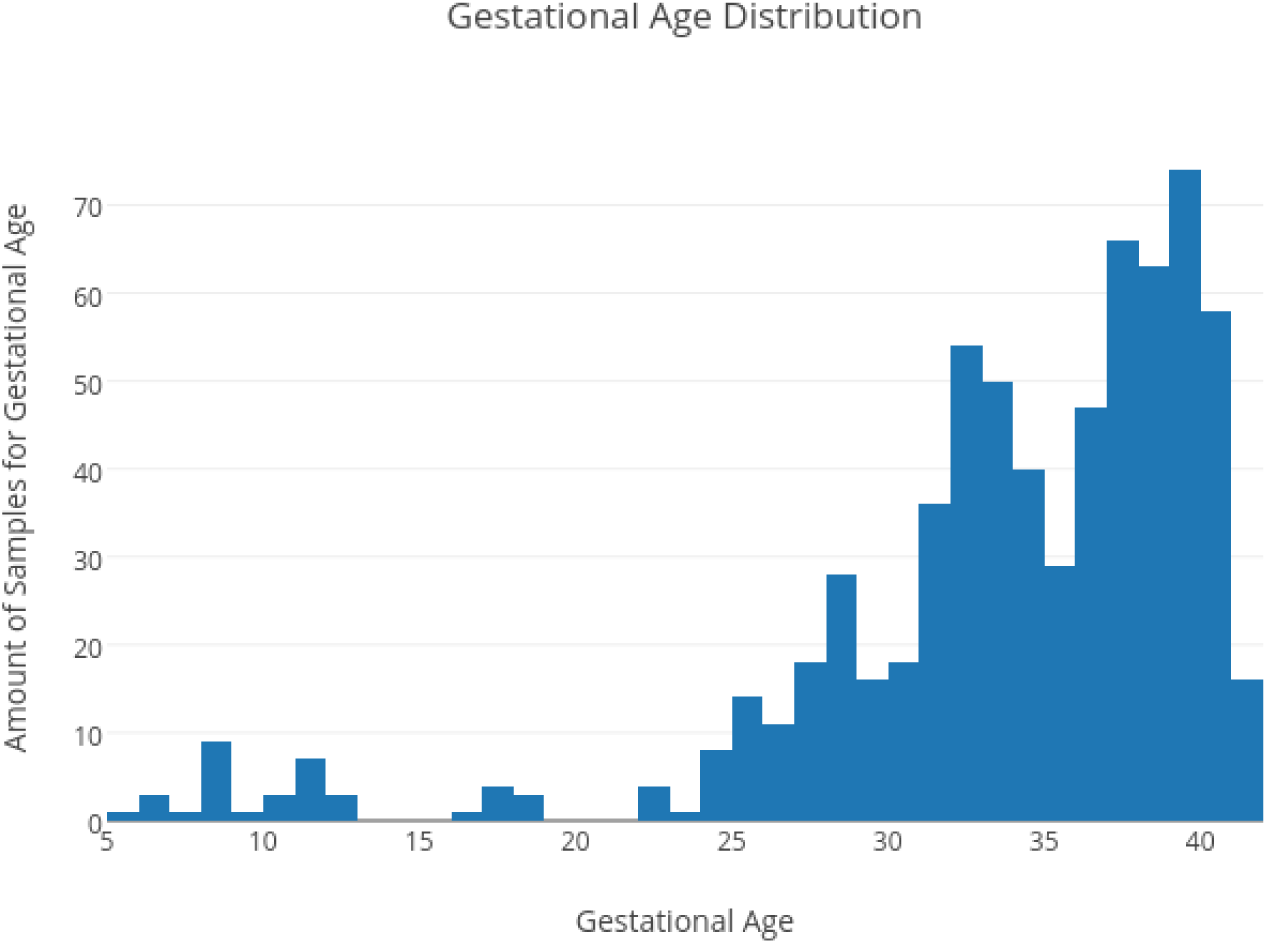
Gestation age distribution for samples with known exact value. Five gestation categories may be considered: first trimester, second trimester early preterm (25 to 30 week), late preterm (30 to 37 week), term (37 to 42 week).

It is worth noticing that the value of gestational age depends on measurement technique. If it is measured as first day of the woman’s last menstrual cycle it is roughly 2 weeks earlier than if measured since actual conception date [6].

Although **Diagnosis** is specified almost everywhere as the majority of the considered experiments are case-control in design, it is not always enough just to know value of this attribute. For example, if the sample is a “control” it can be placenta delivered from natural birth or after c-section, it can be term or preterm birth, medically induced or spontaneous. All these characteristics are to be considered when matching controls to cases during formation of cross-experiment study groups. Likewise, when the sample is “pre-eclampsia” there are also several additional attributes to look at: pre-eclampsia onset, severity, additional complications, formal pre-eclampsia criteria such as blood pressure and urine protein concentration.

**Biological Specimen** is mostly placenta in our study. A biological sample tissue type is specified wherever mentioned to reduce the noise that would be caused by differential expression among cell types inside placenta.

Sample attribute coverage summary for the rest of attributes can be seen in Figure 4. Metadata may be further completed in future either through utilizing existing approximate data as was done here with gestational age or by inferring metadata directly from the expression data as was done, for instance, with fetal sex in [14].

**Figure 4:**
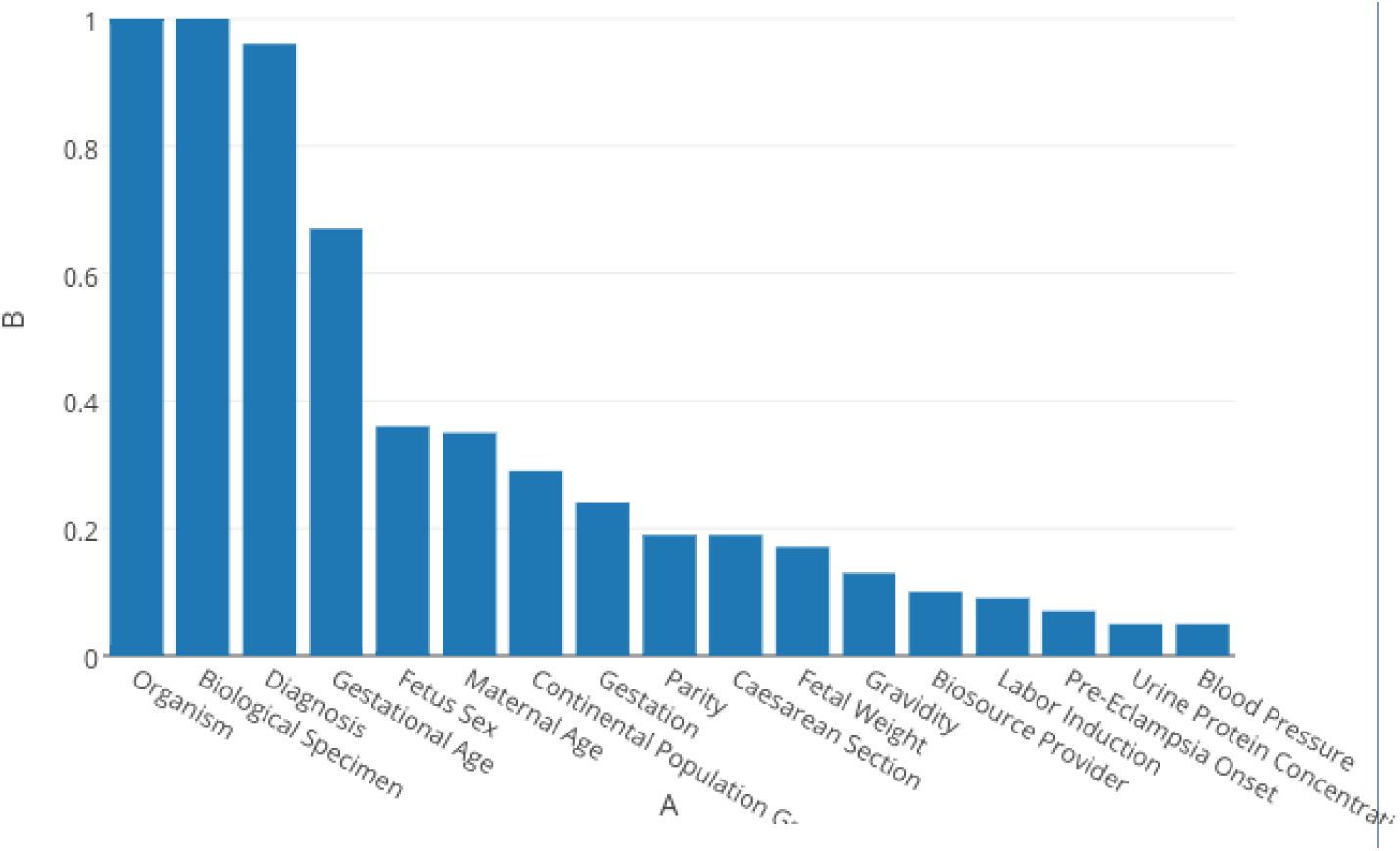
Percentage of samples containing the attribute.

Most of the considered samples are comparable and ready for integrative analysis at this moment. These biological samples can now constitute case-control groups of larger size than in original datasets.

## Discussion

Numerous attempts to organize gene expression data for cross-experiment analysis have been taken by different investigators.

Integrative approach for pre-eclampsia study has been taken in [14], though no database creation was intended. Authors merged 7 gene expression microarray datasets of case-control studies of preeclampsia-affected placenta and created study groups of 77 and 96 samples for pre-eclampsia and control groups respectively. Authors got interesting insights into the nature of the disease that could not have been made from mere observation of individual studies’ results. Namely, unsupervised clustering based on both sample expression and metadata revealed three clusters of samples. One of the clusters turned out to be largely composed of the healthiest term placentas along with supposedly pre-eclamptic ones. A hypothesis was made that unhealthy placentas of that cluster had been misdiagnosed as pre-eclamptic ones, and were really afflicted with another maternal hypertensive disorder, such as gestational hypertension or chronic hypertension.

This is supported by the GSEA and gestational age comparisons, which indicate that cluster 1 is largely composed of the healthiest term placentas in this data set.

**GENEVESTIGATOR** is a search engine for gene expression over a compendium of manually curated datasets [2, 13, 21]. It provides variety of tools addressing the questions of finding conditions under which the genes of interest are the most up- or down-regulated as well as finding the genes that are the most expressed under certain conditions. Biological samples are carefully annotated using custom ontology with a variety of characteristics allowing to filter for highly specific cases. Cross-experiment analysis is based on a concept of meta-profile. As explained in [10] meta-profiles summarize expression levels from many samples according to their biological context. Each sample is annotated with five attributes: anatomical parts, cell lines, cancers types, developmental stages, and perturbations - to generate meta-profiles. Hence each meta-profile is an expression profile of an “average” sample generated from all expression profiles of the samples of specific tissue or developmental stage. An exception is the Perturbation meta-profile, which consists of responses to various experimental conditions (drugs, chemicals, hormones, etc.), diseases, and genotypes. For Perturbation meta-profile results are created by comparing groups of samples from individual experiments. Data from multiple experiments are not mixed to create a single value. As a result, this tool contains large compendia of response types collected from many experiments. It also worth mentioning that samples of two different microarray platforms can not be considered simultaneously.

This implies that GENEVESTIGATOR is closer to meta analysis than to integrative analysis. It integrates sample data for samples of the same platform and biological context to create meta-profiles but other tasks such as comparison of samples of different biological contexts or comparison of differentially expressed genes for different medical or biological cases (perturbations) are pure meta-analysis.

The GENEVESTIGATOR’s does not currently contain datasets of our interest and does non allow user to upload data directly and, thus, can not satisfy our goals.

Another resource enabling interactive query and navigation of transcriptome datasets is **Gene Expression Browser** [18] and its specific implementation for placental gene expression [8]. While its data is more relevant to our pursues and the interface is quite convenient and has some extended features in comparison to GEO or ArrayExpress, such as search for differentially expressed genes in a single experiment, building relevant plots, giving detailed information on found genes and and a bit of meta-analysis tools such as search for experiments that have given gene differentially expressed with a certain rate of difference, no tools for integrative analysis are provided.

An Integrative Meta-Analysis of Expression Data (**INMEX**) service [7] is the one having options for both meta- and integrative analysis, not to mention a huge ensemble of tools for further data analysis. It is worth emphasizing that INMEX is not a database, but a pure tool with no pre-uploaded data except for several examples: uploading raw data and constructing proper study groups are the user’s responsibility. Opportunely, our newly developed database can make for a perfect data input to INMEX and it is quite possible that INMEX’es output will be a solution to our pursue of data integration and subsequent analysis.

Finally, we can not but emphasize how important it is to upload well annotated *sample-wise* experiment metadata. In doing so we join the call from D.M. Nelson and G.J. Burton, editor of the Placenta journal, who published A technical note to improve the reporting of studies of the human placenta featuring those essential parameters attention to which “will lead to enhanced chances that comparisons will be of apples with apples, instead of comparisons of two fruits”[15]. While we notice the improvement in time of the data reporting quality we could not manage to gather even the half of what the note suggests from the publicly available datasets and articles (Figure 4). Namely, the information regarding drugs, previous prenatal admissions, screened for diabetes, antibiotics, beta strep status, antenatal steroids, magnesium sulfate, anesthesia and cervical ripening agent was almost entirely absent.

Despite that, most of the samples in our database are now provided with sufficient metadata for them to be comparable. These biological samples can now constitute case-control groups of larger size than in individual datasets. Described here data gathering and standardization are the first steps to further analysis, namely, the search for differentially expressed genes between different cross-experiment case and control groups. Also, our database is designed and expected to be enlarged with forthcoming gene expression experiments’ metadata and, potentially, with data from other omics (genome, metabolome) with relatively low effort.

## Competing interests

The authors declare that they have no competing interests.

## Author’s contributions

OL, AF designed the database. AF, MO made suggestions on database content. OL implemented database design and its admin web interface and performed metadata standardization. All authors read and approved the final version of the article.

## Acknowledgements

We would like to express out gratitude to Sandra A. Founds for providing clinical data for E-GEOD-12767.

### Additional Files

Supplement 1 — ArrayExpress Experiments’ Accession Numbers

List of ArrayExpress accession numbers for experiments taken for the study.

Supplement 2 — Sample Attribute Names and values

List of all standardized names and values for sample attributes along with the corresponding names and values originally downloaded from ArrayExpress.

